# Understanding interactions between risk factors, and assessing the utility of the additive and multiplicative models through simulations

**DOI:** 10.1101/706234

**Authors:** Lina-Marcela Diaz-Gallo, Boel Brynedal, Helga Westerlind, Rickard Sandberg, Daniel Ramsköld

## Abstract

Interaction analysis is used to investigate the magnitude of an effect which two risk factors have on disease risk, and on each other. To study interactions, both additive and multiplicative models have been used, although their interpretations are not universally understood. In this study, we set out to investigate the resulting interactions of several risk factors relationships in additive or multiplicative scenarios using simulations, in the context of case-control studies. Our simulation set up showed that independent risk factors approach additive relative risk at low disease prevalence. However, risk factors that contribute to the same chain of events (i.e. have synergy) lead to multiplicative relative risk. Additionally, thresholds on the number of required risk factors for a disease (the multifactorial threshold model) lead to intermediaries between additive and multiplicative risks. We proposed a novel measure of interaction consistent with additive, multiplicative and multifactorial threshold models. Finally, we demonstrate the utility of the simulation strategy and discovered relationships on real data by analyzing and interpreting gene-gene odds ratios from a case-control rheumatoid arthritis study.

## INTRODUCTION

Screening the genetics of large cohorts of individuals can identify genetic loci that impact phenotypic traits on a gene-by-gene basis [1], e.g. linking single-nucleotide polymorphisms to traits in genome-wide association studies (GWAS). These studies have identified hundreds of risk genetic variants for large numbers of disorders [2, 3, 4], although such individual genetic associations seldom explain large disease risks alone. It is known that for several diseases combinations of genetic and/or environmental risk factors are important, as they have a larger than expected risk when both factors are present. However, it is still challenging to resolve if, and how, multiple factors interact in shaping traits, and to biologically interpret the identified interactions [5].

The association between individual genetic loci and an outcome (e.g. disease) is typically quantified as odds ratios or relative risks. Often the case-control design is used to query low prevalence diseases, in which odds ratios approximate the relative risk in the population even though the samples are unevenly drawn. Interaction tests among risk factors are often examined pairwise, yielding three odds (or risk) ratios notated as [6]: OR_11_ for carrying both risk factors; OR_10_ and OR_01_ for the exclusive combinations. The lack of both risk factors OR_00_, is used as reference (OR_00_=1). Two different models are commonly used to test interaction, the additive (OR_11_ = OR_10_ + OR_01_ – 1) and the multiplicative (OR_11_ = OR_10_ · OR_01_). The additive model builds on work by KJ Rothman [7], who showed that if two factors are part of the disease’s cause and are part of the same sufficient cause (e.g. pathway), then their joint risks will be larger than their sum (often termed “departure from additivity”). This additive model has been criticized for always giving positive results [8, 9]. On the other hand, the multiplicative model has been criticized as a statistical convenience without a theoretical basis, boosted by the implicit multiplicativity in logistic regression [8, 10, 11]. There is often confusion regarding when and how to use and interpret each of the mentioned models.

In this study we strive to lessen the confusion by using simulations of various interaction scenarios. We thereafter show how additive and multiplicative risk scales compare to our simulation scenarios, aiming to contribute to understand and interpret different models of interaction. Simulations are able to give an answer with high precision level if enough data points are generated. We designed a variety of different scenarios as basis for the simulations. Our intension was to include the most likely real scenarios. In some scenarios we give the cause of the outcome (disease or otherwise) in its description. Others model confounder situations, where there is no cause in the description yet there are apparent, false, risk factors. We will later show how the modelled scenarios can be distinguished in real data, and where they cannot. Across all simulations the risk factors are dichotomous, and neither necessary nor sufficient for disease to occur. All but one simulated scenarios include two risk factors. Limited statistical power restricts the possibility to test for more than two factor interactions in real data, and as the three risk factor scenario will illustrate, it is possible to elucidate multifactorial interactions from the set of pairwise interactions.

## METHODS

### Simulations

To better understand risk interactions, we first set up simulations of causation, which are represented in Figures 1A to 1C. Each scenario included a dichotomous outcome (present/absent), and causative processes termed components. One thousand simulations were performed for each of the different simulation scenarios (Figs 1A-C, 2B, 3A-B, S1A-C and S2A-B). Each simulation consisted of one million data points (or simulated individuals) where the presence or absence of a risk allele was assigned, as well as the status of case or control according to each model. Since simulations with 100,000 data points per simulation gave essentially the same results, these ones were not included here. For the sake of simplicity, binary factors were used, corresponding to a dominant or recessive scenario in genetics. We used subgroups with the same ratio of cases to controls, named components, except for the three factors model (Fig 2B), where components 2 and 3 had the same ratio and component 1 has the same ratio as the logical OR combination as those two.

**Figure 1.**
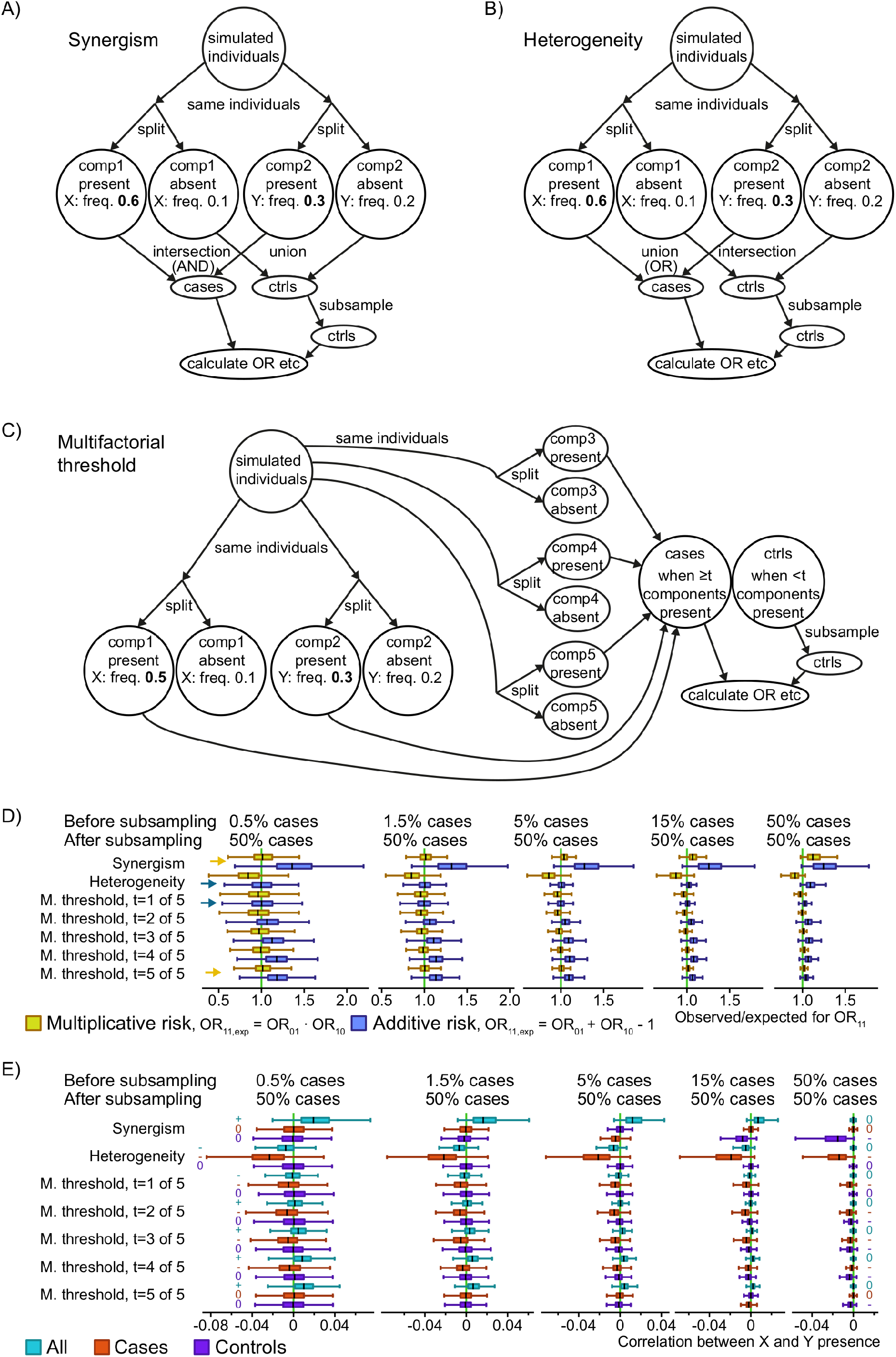
Three simulated causal scenarios with selection of equal numbers of cases and controls. A-C) Simulation schemes for three generalized scenarios in a case-control study context: synergism of causes (A), heterogeneity of causes (B) and a multifactorial or 5-factor threshold (C). The numbers are example frequencies, and frequencies in bold highlight the higher frequencies of the simulated risk factors (X and Y) associated with disease. For example, “Y: freq. 0.3” means that each simulated individual in the group had a 30% chance of being assigned the risk factor Y. The numbers in italics are the average frequency in the other group of simulated individuals, note that this will depend on the prevalence (which is adjusted in the scenarios in the “split” into cases and controls). “Components” (comp1 and comp2) were used as a strategy to obtain probabilistic risk factors. D) The odds ratios for double risk (OR_11_) calculated from the simulation scenarios, with boxes summarizing 1,000 simulation runs with different risk factor frequencies. The observed OR_11_ were compared to the additive and multiplicative combinations of the odds ratios for single risk (OR_10_ and OR_01_). Boxplots show median and quartiles for the simulations, but extreme values are omitted for clarity. Yellow arrows highlight where the median is visibly close to multiplicativity, while blue arrows do the same for additivity, for the two most extreme simulated prevalence rates. “M. threshold” refers to multifactorial threshold (scenario C). E) Correlation coefficients between the risk factors X and Y, for three sample sets (all, cases only, controls only). The relevant signal in each case is whether the median is negative, zero or positive, highlighted with a -, 0 or + symbol for the two most extreme simulated prevalence rates.

**Fig 2.**
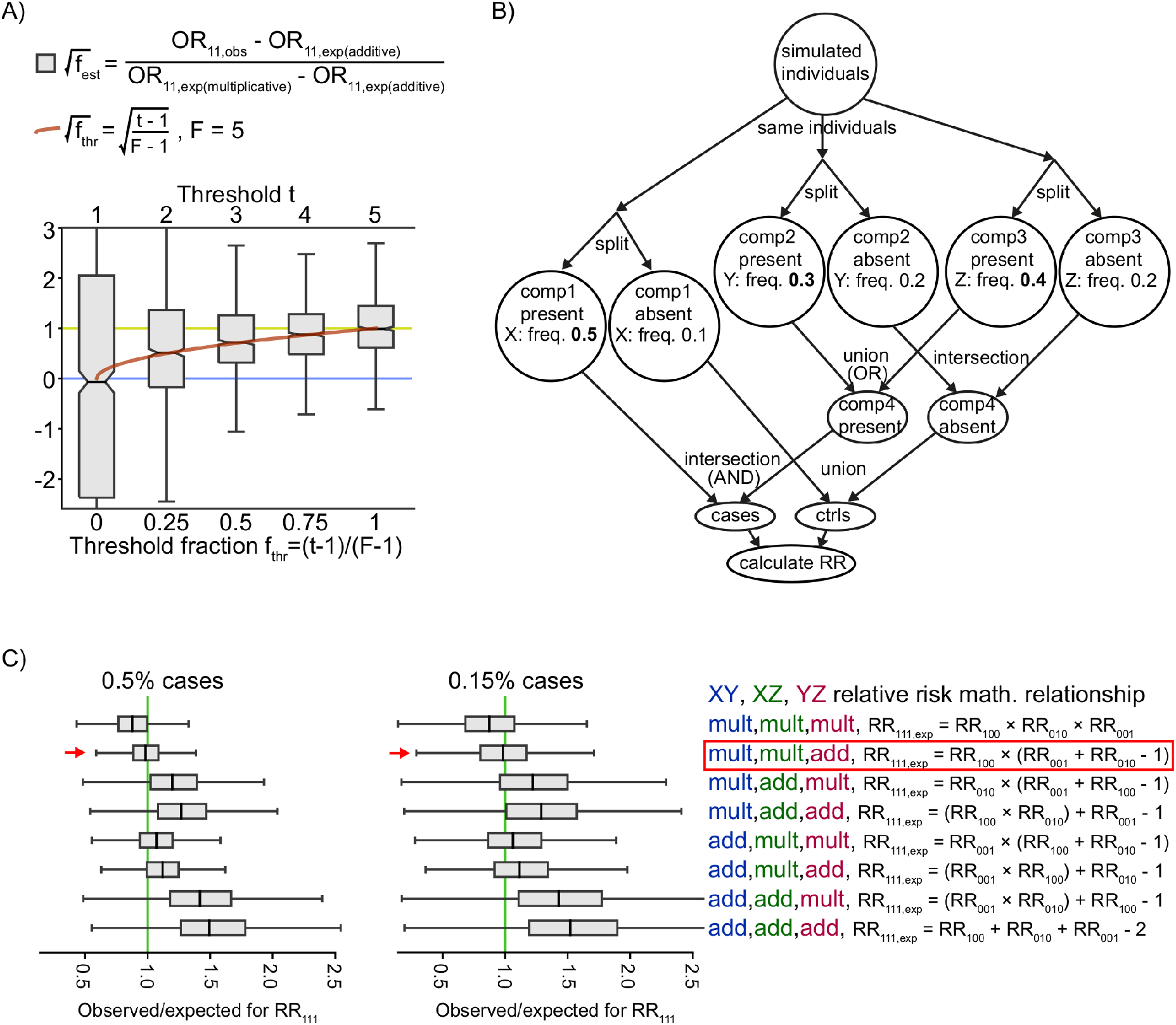
Association type estimation of the multifactorial threshold model and a 3-factor causal scenario. A) Results using the scenario in Fig 1C. The red curve is the square root of the threshold fraction; this and √f_est_ make up the y axis. Notched in the box plots show bootstrapped 95% confidence intervals for the medians. The simulation had 0.5% cases and the rest controls, with a range ±0.04%. B) The numbers are example frequencies, and frequencies in bold highlight the higher frequencies representing association with disease. For example, “Y: freq. 0.3” means that each simulated individual in the group had a 30% chance of being assigned the risk factor Y. This scenario required one “component” (where X is associated with risk) together with either of two components (where either Y or Z is associated with risk) to produce the outcome. C) The observed relative risk from the simulation in (B) compared with the 8 possible combinations of additive (add) and multiplicative (mult) combinations. “mult, mult, add” is marked by a box, as (B) should theoretically give an AND relationship between X and Y and between X and Z but an OR relationship between Y and Z. Therefore, RR_100_ (for X alone) is multiplied by the additive for Y and Z’s relative risks, assuming logical AND give multiplicative while logical OR give additive relative risks and odds ratios at low prevalence.

**Fig 3.**
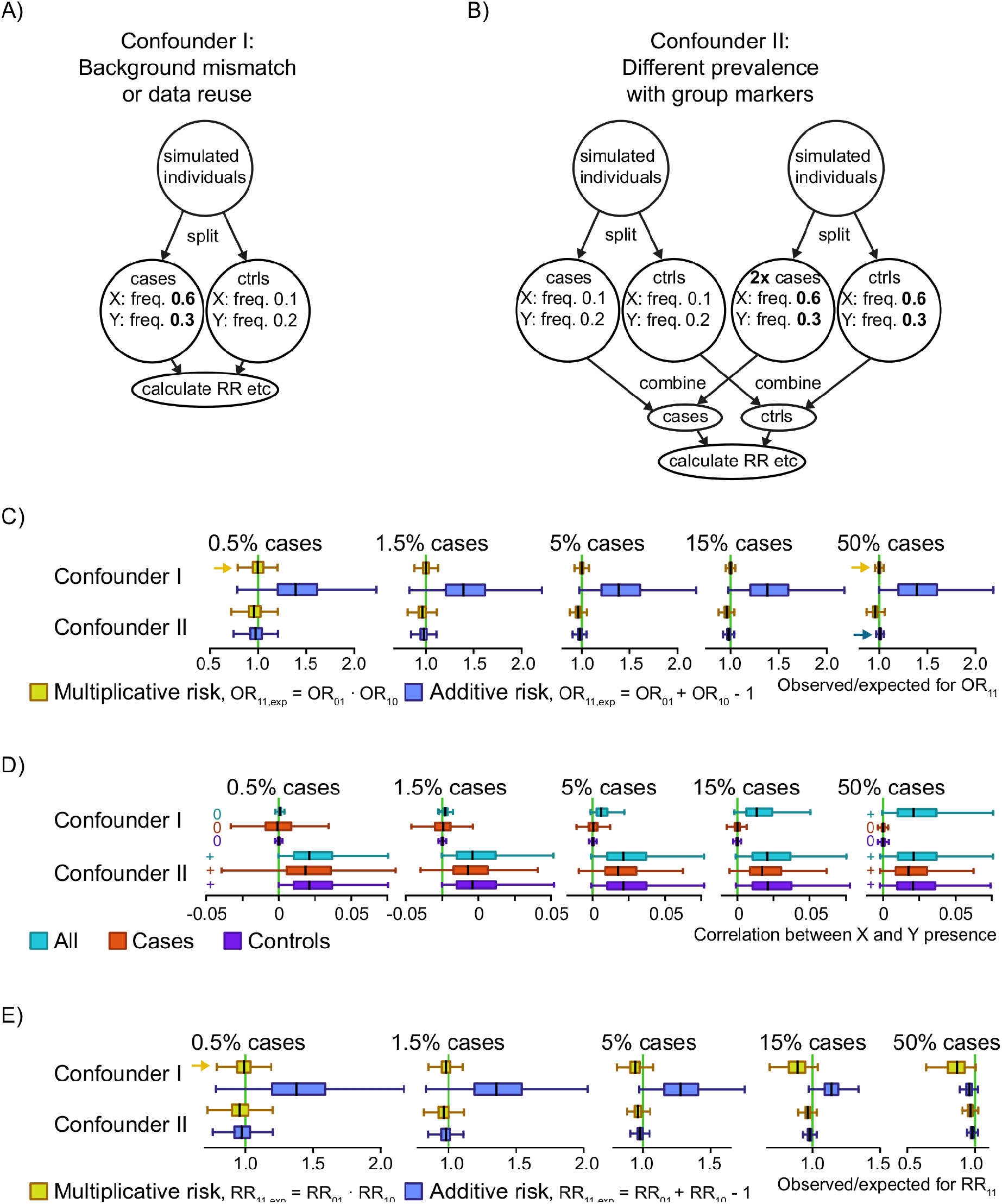
Two simulated confounder scenarios. A-C) Simulation schemes for two generalized scenarios: either background mismatch between cases and controls or testing interactions on the same data as selecting risk factors (A), and a mix of groups with both different background and different prevalence rates (B). The numbers are example frequencies, and frequencies in bold highlight the higher frequencies of the simulated risk factors (X and Y) associated with disease. For example, “Y: freq. 0.3” means that each simulated individual in the group had a 30% chance of being assigned the risk factor Y. In each indicated circle the two risk factors have been added in an uncorrelated/independent manner (C) The odds ratios for double risk (OR_11_) calculated from the simulations, with boxes summarizing 1,000 simulation runs with different risk factor frequencies. The observed OR_11_ were compared to the additive and multiplicative combinations of the relative risks for single risk (OR_10_ and OR_01_). Boxplots show median and quartiles for the simulations, but extreme values are omitted for clarity. Yellow arrows highlight where the median is visibly close to multiplicativity, while blue arrows do the same for additivity, for the two most extreme simulated prevalence rates. D) Correlation coefficients between the risk factors X and Y. The relevant signal in each case is whether the median is negative, zero or positive, highlighted with a -, 0 or + symbol for the two most extreme simulated prevalence rates. E) Like (C) but for relative risk.

The allele frequency for the two exposures tested in each model, named X and Y, was established before each simulation and set from a random value in a given range. For instance, the lower frequency for the risk factor Y was a random value between 5% to 15%. Then the higher frequency of the factor Y was the multiplication of the lower frequency by a random number between 1.1 to 4. Similarly, the lower frequency for the factor X was randomly designated between 5% to 25%. In turn, the higher frequency of the factor X was the multiplication of its lower frequency by a random number between 1.1 to 2. The status of the risk factors in the simulations were set randomly in accordance with the frequencies, so that e.g. a frequency of 0.2 meant of 20% chance of being assigned a 1 and 80% to be assigned a 0 for that risk factor.

For each scenario we calculated the interactions between two risk factors. Due to the nature of the components, more risk factors can exist in each scenario without interfering with the calculations. For each component there was one risk factor that was explicitly made to correlate with its “present” state (Fig 1A-C), and the risk factor frequencies were varied between simulation runs. We subsampled the controls to match the number of cases at the end, to resemble how case-control studies reflect the population in a biased way (Fig 1). We calculate several metrics for each scenario. For the scenarios with explicit cause and biased subsampling we calculate odds ratios for the combinations of the risk factors (OR_01_ for *X*=0, *Y*=1, OR_10_ for *X*=1, *Y*=0, OR_11_ for *X*=1, *Y*=1; Fig 1D) and Pearson correlation coefficients between the risk factors X and Y (Fig 1E). For the versions of the scenarios without the subsampling (S1 Fig) we calculate relative risk instead, as that is the appropriate population measure.

### Interactions in rheumatoid arthritis GWAS

We evaluated both additive and multiplicative interaction on a human case-control genome-wide association dataset for anti-citrullinated protein antibody positive (ACPA-positive) rheumatoid arthritis (RA), from the Swedish epidemiological investigation of RA (EIRA) cohort. For the two top genetic risk factors for RA in European-descendent populations, *HLA-DRB1* shared epitope and *PTPN22* rs2476601 T, we tested these risk factors against all non-HLA risk SNPs. *HLA-DRB1* shared epitope is a group of alleles with similar effect, and rs2476601 is a non-synonymous coding variant of the *PTPN22* gene. Two tests were used, one which used additivity as hypothesis and other that used multiplicativity as hypothesis.

The genotyped and imputed GWAS data from the EIRA study were used in this part of the study (see [12] for sources included). Only data from ACPA-positive patients with RA was included. The standard data filtering was performed as previously described [12]. Briefly, genotyping missing rate higher or equal to 5% and *P*-values of less than 0.001 for Hardy-Weinberg equilibrium in controls were excluded. The SNPs located in the extended HLA region (chr6:27339429-34586722, GRCh37/hg19) were removed, due to the high linkage disequilibrium and possible independent signals of association with ACPA-positive RA in the locus.

The departure from additivity or multiplicativity for two risk factors was estimated in the imputed GWAS data (3,138,911 SNPs for the test with *HLA-DRB1* shared epitope and 3,308,784 SNPs for the test with *PTPN22* rs2476601 T), using GEISA [13], where a dominant model was assumed. The first ten principal components and gender were used as covariates in this analysis, in order to control by population stratification and differences between allele frequencies due to sex, respectively. A cut-off of minimum five individuals per each odds ratio combination was applied. The *HLA-DRB1* shared epitope alleles included *01 (except *0103), *0404, *0405 and *0408 and *1001. The *P*-values for interaction from these analyses are plotted in the Fig 4A-B. We included only SNPs at risk allele frequencies between 10% and 50% in this testing to minimize the risk of including protective factors, however when we tested all the SNPs at a minor allele frequency above 1% the result provided the same conclusion.

**Fig 4.**
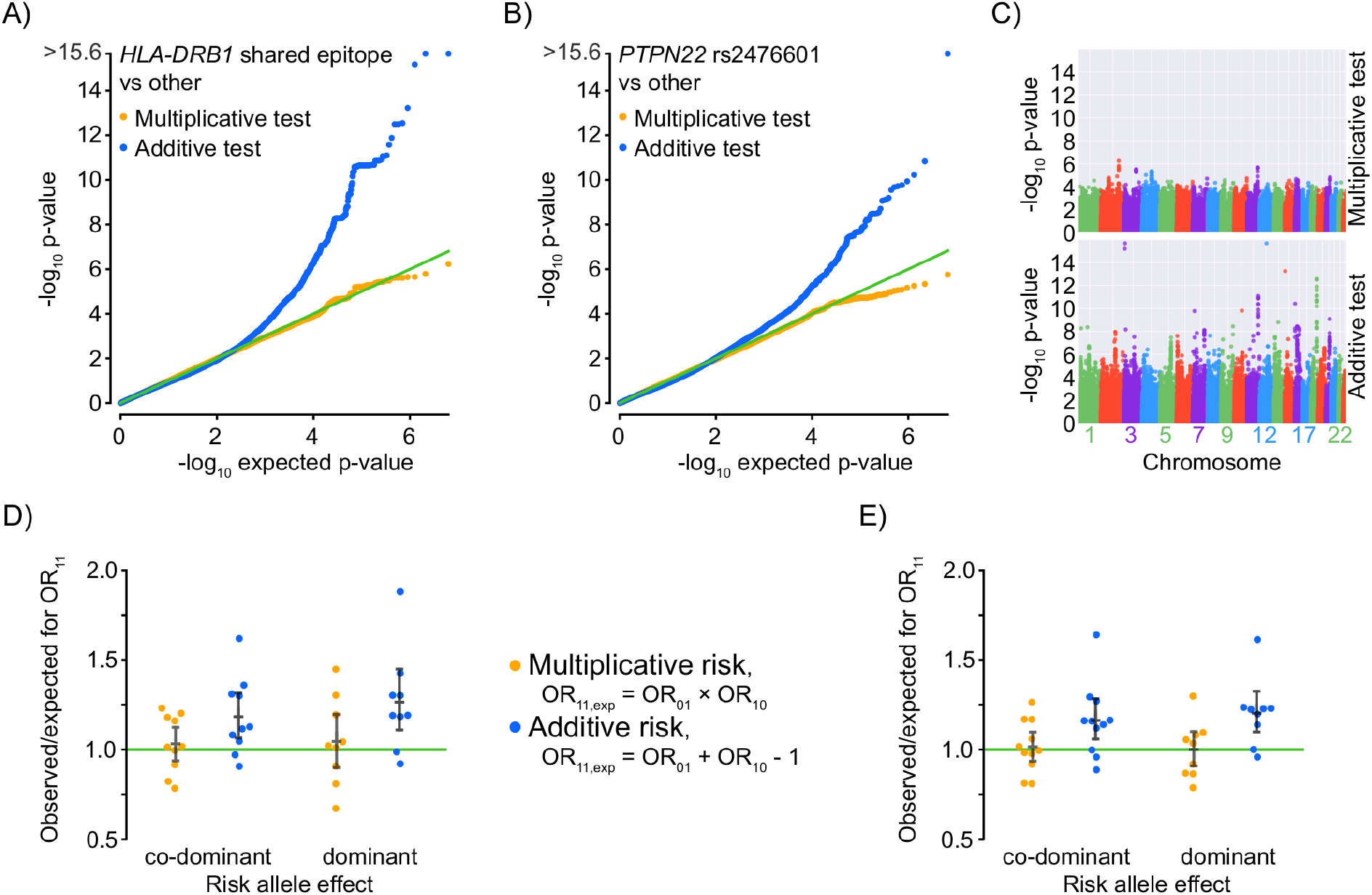
Application to genome-wide association data for rheumatoid arthritis. A-B) *P*-value distribution for two tests, one for deviation from additivity and one from multiplicativity. Two risk factors (see text for alleles) are tested in EIRA data against the rest of the genome except nearby SNPs. Each bin is 0.01 wide. A uniform distribution means a lack of deviation from the null model. C) Manhattan plots for deviation from multiplicativity or additivity, for *HLA-DRB1* shared epitope against the rest of the genome except nearby SNPs. D) Odds ratio for *HLA-DRB1* shared epitope and one other risk factor, compared to the expected from an additive or multiplicative null model. Only known RA risk SNPs from the literature are shown. Black bars show median and 95% confidence intervals (bootstrap). E) The same SNPs as for (D) but shuffled within cases and shuffled within controls to match Confounder I, thereby being a positive control for multiplicative odds ratios, allowing a comparison of dispersion with (D).

To address both additive and multiplicative risk scales and evaluate the behavior of the odd ratios for double risk exposure (OR_11_ – Figs 4C-D, S3 and S4), we used genotyped EIRA GWAS data (281,195 SNPs). For this analysis, the data was transposed using Plink 1.07 [14]. Known risk SNPs were selected based on an odds ratio higher than 1.1 and 95% confidence intervals do not overlapping 1, together with the criterion of having been reported as associated to RA in published case-control RA GWAS [15, 16, 17].

### Computational packages

Calculations were done using python, including the packages numpy [18], scipy [19], matplotlib [20], pandas [21], scikit-learn [22], seaborn [23], jupyter and geneview [24]. Pearson correlation was used to calculate correlation coefficients.

### Data access

Code is available at https://github.com/danielramskold/additive_risk_heterogeneity_multiplicative_risk_synergism2 where we also provide the code used to generate the figures.

## RESULTS

### Scenarios with risk factors contributing to the explicitly causal components

In the synergism scenario, two components are needed to be in the state “present” for the outcome “case” to occur (Boolean AND logic; Fig 1A). In the heterogeneity scenario, it is enough that either one of two components are in the “present” state for a “case” outcome (Boolean OR logic; Fig 1B). We also sought to investigate the multifactorial threshold model, which has been thought of in terms of genetic liability [25]. Here we chose five components, although any number from three could have been used. In this model a minimum number of factors need to be present for causation, which translated to our simulation mean that at least *t* number of components need to be in the “present” state for a “case” outcome (Fig 1C). We also present the results without subsampling, which should match population studies (S1 Fig). For the two risk factors, the synergism scenario corresponds to Rothman’s synergism concept [26], and the heterogeneity scenario corresponds to genetic heterogeneity.

For phenotypes with a low prevalence the presence of both risk factors had a multiplicative association in the synergism scenario (Fig 1D). For the heterogeneity scenario the presence of both risk factors instead had an additive association low prevalence (Fig 1D). The multifactorial threshold scenario imitated the heterogeneity scenario at threshold *t=1* and the synergism scenario at *t*=5 (where 5 meant *all* components needed to be present for phenotype to occur). For intermediate thresholds in the multifactorial threshold model we obtained intermediate results so that at low prevalence the joint behavior of the factors was above additive but below multiplicative (Fig 1D). Without the subsampling (S1E Fig), the scenarios which produced multiplicative associations lacked correlation between risk factors, both within cases, controls, and the two combined (“all”). However, the subsampling caused a positive correlation between risk factors in the combined (“all”) group (Fig 1E). The additive and intermediate associations on the other hand, without the subsampling, were reflected in negative correlation within cases, yet still not in controls (S1E Fig). This negative correlation among cases for two risk factors for additive models can be understood theoretically, since two individually sufficient factors (meaning that there is heterogeneity of causation) should have a strongly negative correlation. A similar, but attenuated, pattern for non-sufficient risk factors in regards to correlation is thus not surprising. We also present a simulation of population (or unbiased random sampling) results, i.e. without the subsampling step, and there we found a multiplicative association on relative risk in synergism scenario between the risk factors at every prevalence rate (fraction of cases in the population), whereas for the heterogeneity scenario the association was additive at low prevalence but less-than-additive at high prevalence.

### An interaction metric with a simple interpretation for multifactorial thresholds

As we have seen, a risk factor relationship in the multifactorial threshold model gives rise to intermediary interaction terms between additivity and multiplicativity (Fig 1D). We therefore wanted to find a formula describing *how* intermediary is. We scaled the threshold as the ratio *f*_thr_ = (*t* – 1) / (*F* – 1) where *F* is the number of components (factors) that could cross the threshold *t*, in order to have a 0 to 1 scale, and we scaled the odds ratio between additive, at 0, and multiplicative, at 1 (see Fig 2). We found that the square of the threshold ratio fit the scaled odds ratio. This means that the odds ratio at a low prevalence (such as 0.5%) for doubly exposed was (1 – √*f*) · expected(additive) + √*f* · expected(multiplicative), i.e. OR_11_ = OR_10_ + OR_01_ – 1 + (OR_10_ – 1) · (OR_01_ – 1) · √*f*, where √*f* is the square root of *f*_thr_. The metric used, √*f*_est_ (signed square root of the estimated multifactorial threshold fraction), has the convenient properties of having both a natural lower value (0 for additive) and higher value (1 for multiplicative) as well as an interpretable scale in-between through its connection to the threshold fraction *f*_thr_. While the metric *f*_est_ = sign(√*f*_est_) · (√*f*_est_)^2^ had a more natural scale, unlike √f_est_ this was not symmetric around the median (S2 Fig). The large spread for √*f*_est_ at threshold 1 (Fig 3C) could have resulted from having cases that did not involve components 1 or 2 in this simulation scenario (Fig 3A) and thus lowering odds ratios, rather than intrinsically from additive risk. The metric √*f*_est_ is related to another measure of interaction size, relative excess risk due to interaction (RERI), by √*f*_est_ = RERI / (OR_10_ – 1) / (OR_01_ – 1) and could perhaps be used instead of measures like attributable proportion, synergy index and RERI, given that it can pinpoint multiplicative risk in testing on the additive scale and vice versa.

### Three risk factors

As could be seen in our simulation scenario based on the multifactorial threshold model, even relationships involving more than two potential causes can be analyzed by pair-wise interactions. But an outcome with more than two causes does not entail a threshold model. It should also be possible to break down into pairwise risk factors relationships those involving causal heterogeneity and synergism. For example, in the case of outcome = *X* AND (*Y* OR *Z*) = (*X* AND *Y*) OR (*X* AND *Z*), the risk factors *X, Y* and *Z* the relationship should be distinguishable as a heterogeneic between *Y* and *Z* (from the logical OR) and synergism between *X* and each of the other risk factors (from the logical AND) (Fig 2B). We tested this example against the expected relative risk from all eight possible combinations of logical OR and AND, and found the simulation to perform as predicted (Fig 2C, S1C Fig).

### Scenarios where confounder effects cause false interaction

We investigated two confounder scenarios, neither of which include any causal relationship between the risk factors (e.g. heterogeneity or synergism) and are therefore most applicable when either or both are false risk factors. The first describes when the genetic background for cases and controls are mismatched (Fig 3A). It can also describe an effect of data reuse, where the same data set is used to define risk factors and to calculate interactions [9]. The other scenario we simulated is when there are multiple groups with different markers frequencies and different prevalences between groups (population stratification), in our case we condensing it down to two groups (Fig 3B). Our simulations could not model linkage disequilibrium (LD), but risk factors pairs in LD are regularly filtered out in interaction analyses.

For the first confounder scenario (Fig 3A), odds ratios (Fig 3C) and correlations (Fig 3D) at high fractions of cases are more relevant than relative risks (Fig 3E) at fractions of cases corresponding to disease-level prevalence. Since the associations are not generated by the population structure itself and should therefore instead mirror study design. We found the odds ratio to be multiplicative, regardless of the balance between cases and controls. This makes it indistinguishable from synergism in terms of association. This was also true for correlation coefficients, where only population data would be able to distinguish synergism from this confounder scenario (Fig 1E, S1E Fig, Fig 3D) by the former’s lack of correlation in that situation. The second confounder scenario (Fig 3B) had below-additive odds ratio and relative risk, matching additive at a high fraction of cases (Fig 3C, 3E). The risk factors correlation among the three groups used (all, cases, controls) was always positive (Fig 3D), which should be useful for distinguishing this scenario from others. We also tested what happens when adding a biased subsampling step as in Fig 1A-C to Confounder II but it made no difference to the results (data not shown). Out of interest for additive associations, and because such schemes could guide randomization, we also devised one simulation scheme that always produced additive relative risk and one that always produced simulation scheme additive odds ratios (S3 Fig), complementing the always-multiplicative odds ratio of the first confounder scenario.

### Example from rheumatoid arthritis

Both the synergism and heterogeneity scenarios represent interesting relationships between two risk factors, thus the appropriate interaction model to use depends on the hypothesis one is interested in. Given the lack of insight into many RA risk loci, we needed a hypothesis-free approach, and the closest to that is evaluating both additive and multiplicative hypotheses, as that would cover both models as well as threshold-based scenarios due to their intermediate nature (i.e. they would fail both types of tests in the opposite direction). We therefore evaluated both additive and multiplicative interaction for the two top genetic risk factors for RA in European-descendent populations, *HLA-DRB1* shared epitope and *PTPN22* rs2476601 T allele, on GWAS data from the EIRA study. We detected no deviation for multiplicativity, but (as reported before for *HLA-DRB1* [12]) from additivity (Fig 4A-B). The new simulation presented here increases our ability to interpret this result as a widespread interaction between *HLA-DRB1* shared epitope and all non-*HLA* genetic risk factors, in the common meaning of interaction where synergism is a type of interaction. Correlation analyses backed up synergism (but not heterogeneity or population stratification) as match for the results (S4 Fig). From this, we could derive that the *HLA-DRB1* shared epitope cannot be substituted for (i.e. phenocopied by) a non-*HLA* genetic risk factor for its part in the chain of ACPA-positive RA etiology (Fig 4A,C,D). The same was the case for the *PTPN22* risk allele, given the similarities in *P*-value distributions we observed (Fig 4A-B). For both set of tests there were a majority of tested loci where there was too little data to distinguish additive from multiplicative odds ratios. We followed up the results of multiplicativity by looking only at known risk SNPs (Fig 4D), finding results similar to a randomization based on the Confounder I scenario (and therefore bound to produce multiplicative odds ratios), with similar variability (*P*=0.6-0.8, Levene’s test, *n*=9-11 SNPs) implying a dearth of non-multiplicative odds ratios (Fig 4E). This randomization is the same as Test III of [27]. We also devised a randomization scheme creating additive odds ratios based on the Additive O scheme of S3 Fig and tested it on those full SNP set (S5 Fig), where it as expected deviated very noticeably from the real data.

## DISCUSSION

We herein present a simulation approach intended to help interpretation of additive and multiplicative interaction of relative risks and odds ratios. We show that additivity of odds ratios occurs when the risk is comprised of two independent risk factors (heterogeneity scenario, Fig 1B), or a process that approximates that setup at a given fraction of cases (S3 Fig). Multiplicativity of odds ratios, and deviation from additivity, follows if two different mechanisms are required for disease (synergism scenario, Fig 1A). Depending on the hypothesis one should therefore chose the appropriate statistical test. To find deviation from synergism, this could be tested using deviation from a multiplicative association. Often however, we are interested in testing whether disease is caused by the interaction of two factors, and then it is appropriate to test for deviation from additivity.

A multiplicative assumption would have merit in our testing against *HLA-DRB1* shared epitope and *PTPN22*, if ACPA-positive RA were a homogeneous set of causes, rather than the kind of heterogeneity of causation that we have shown give rise to additivity between risk factors. Despite being defined based on a mediating risk factor, such homogeneity of ACPA-positive RA is not thought to be the case [28]. But in this paper, we did also test the multiplicative interaction between the strongest genetic risk for ACPA-positive RA and other risk-SNPs in the same material as [12], and showed that it always conforms with the multiplicative null (admittedly, we only tested the main other model – not for example a threshold model with few absent factors allowed). In light of the new understanding that our simulations give, the presence of deviation from additivity, along with no deviation from multiplicativity, supports the existence of widespread synergism between the genetic risk factors in causing ACPA-positive RA. The result is also applicable to the calculation of heritability for RA, since the assumption of an additive model would lead to an underestimation of narrow-sense heritability [29].

Synergism can be viewed as the chain of events scenario, whereas causal heterogeneity corresponds to phenocopying. In terms of Rothman’s sufficient-cause model, the risk factors X and Y in the synergism scenario correspond to risk factors in the same cause, referred to as causal co-action, joint action or synergism, whereas X and Y in the heterogeneity scenario correspond to risk factors in different causes [26].

The fact that most loci showed no statistically significant deviation from neither additive nor multiplicative interaction will be the unfortunate reality for many applications of interaction testing. While statistical power for single risk factor testing scales with the inverse square of the number of samples, already forcing large GWAS sample sizes, the statistical power for interaction testing scales to the inverse power of four [29], thus requiring far larger sample sizes than standard association testing.

Our inspiration for this work came from a simulation study [9] in which case or control status was randomly assigned, and one risk factor was simulated to resemble the strongest genetic risk factor for ACPA-positive RA, and interaction with other risk factors (selected by *P*-value for risk) was computed. The simulation led to an overrepresentation of additive interactions (i.e. deviation for additive odds ratios). Our Confounder I scenario (Fig 3A) was produced an analogous result. The author [9] noted that his simulation produced a multiplicative null model that does not match additivity and concluded that the additive interaction observed were erroneous, as no interaction should be present. We propose an alternative interpretation, based on the convergences we found at low prevalence: the Confounder I scenario has an intrinsic interaction in the meaning of synergy between the risk factors [30] as the model is equivalent to taking the synergism scenario at low prevalence and subsampling it with a bias for cases (unchanged odds ratio means Confounder I is unaffected by biased sampling, and the equal match to multiplicative model at low prevalence, and in terms of correlations, means they become the same at a prevalence like that of RA: 0.7% for all RA in Sweden [31], of which 60% are ACPA-positive [32]). The simulation study [9] is therefore in line with the confusion over additive and multiplicative interaction that can sometimes be found in the literature [33], highlighting the need to understand the relationships to the risk factors that they imply.

In this simulation study we demonstrate the causal interpretations of additive and multiplicative interaction in both the relative risk and odds ratio settings. Some of this has been understood intuitively in the past, especially the connection between multiplicative effect and logical AND [34], but here we try to further show this to the reader through simulation. We hope that this will guide the interpretation of future interaction studies.

## ACKNOWLEDGEMENTS

Anton Larsson helped during literature search. Lars Klareskog, Lars Alfredsson and Leonid Padyukov engaged in scientific discussions. This research was funded in part by Ulla och Gustaf af Uggla Foundation (2018-02670), Reumatikerförbundet (R-861801) and Konung Gustaf V:s 80-årsfond (FAI-2018-0518).

## AUTHOR CONTRIBUTIONS

Conceptualization: D.R. and B.B.; Methodology: D.R. and H.W.; Software: D.R.; Formal Analysis: D.R. and L.M.D.G.; Resources: L.M.D.G.; Writing – Original Draft Preparation: D.R.; Writing – Review & Editing: B.B., R.S., D.R., L.M.D.G. and H.W.; Visualization: D.R.; Supervision: L.M.D.G. and R.S.

## ABBREVIATIONS

ACPA: Anti–citrullinated protein antibody
EIRA: Epidemiological investigation of rheumatoid arthritis
GWAS: Genome-wide association study
HLA: Human leukocyte antigen
OR: Odds ratio
RA: Rheumatoid arthritis
RERI: Relative excess risk due to interaction
RR: Relative risk
SNP: Single-nucleotide polymorphism

