## Supplemental Figures for "Understanding interactions between risk factors, and assessing the utility of the additive and multiplicative models through simulations"

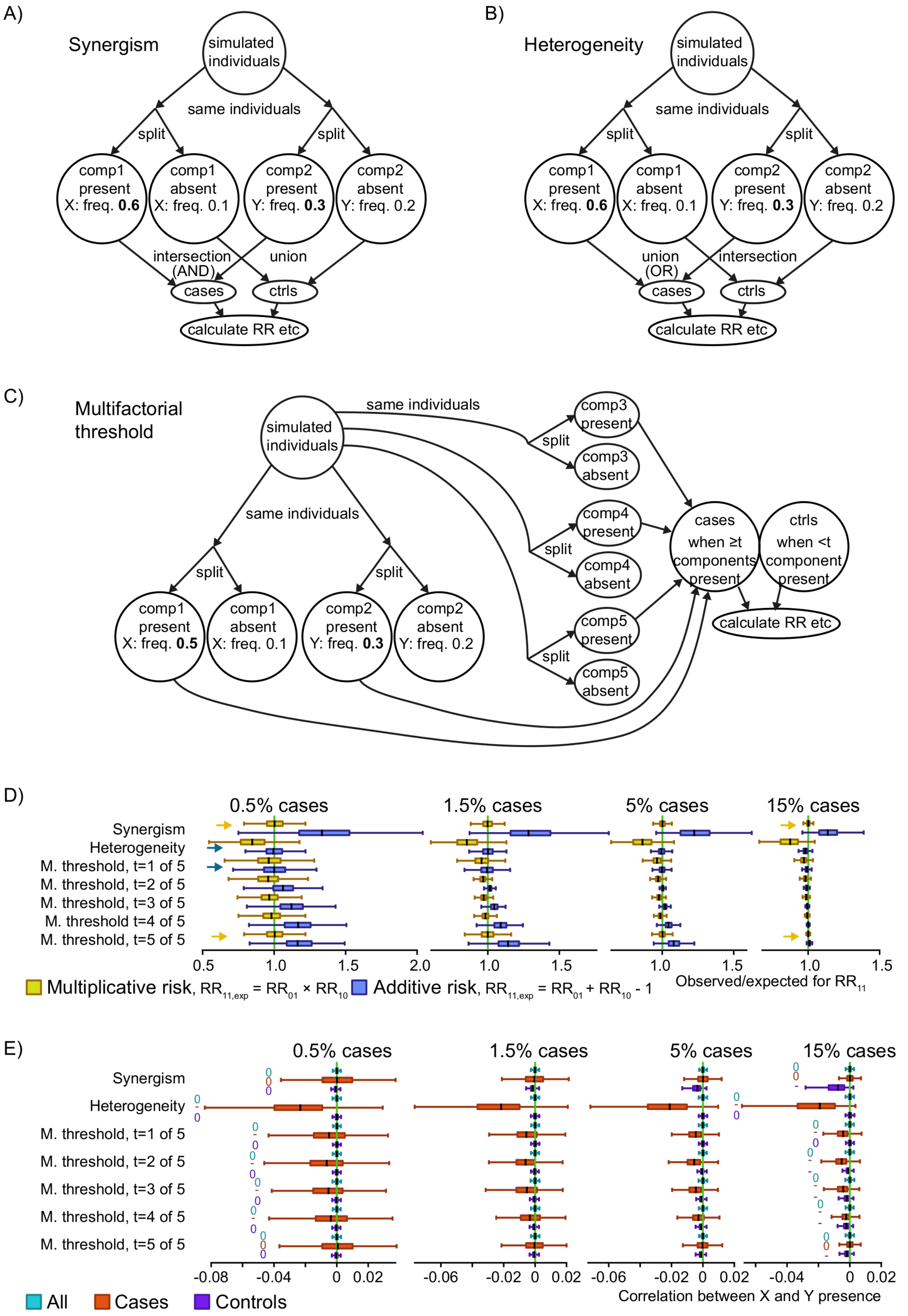

**S1 Fig. Three simulated causal scenarios.** A-C) Simulation schemes for three generalized scenarios: synergism of causes (A), heterogeneity of causes (B) and a 5-factor threshold (C). The numbers are example frequencies, and frequencies in bold highlight the higher frequencies of the simulated risk factors (X and Y) associated with disease. For example, "Y: freq. 0.3" means that each simulated individual in the group had a 30% chance of being assigned the risk factor Y. The numbers in italics are the average frequency in the other group of simulated individuals, note that this will depend on the prevalence (which is adjusted in the scenarios in the split to cases and controls). "Components" (comp1 and comp2) were used as a strategy to obtain probabilistic risk factors. D) The relative risks for double risk ($RR_{11}$) calculated from the simulation scenarios, with boxes summarizing 1,000 simulation runs with different risk factor frequencies. The observed $RR_{11}$ were compared to the additive and multiplicative combinations of the relative risks for single risk ($RR_{10}$ and $RR_{01}$). Boxplots show median and quartiles for the simulations, but extreme values are omitted for clarity. Yellow arrows highlight where the median is visibly close to multiplicativity, while blue arrows do the same for additivity, for the two most extreme simulated prevalence rates. "M. threshold" means scenario C. E) Correlation coefficients between the risk factors X and Y. The relevant signal in each case is whether the median is negative, zero or positive, highlighted with a -, 0 or + symbol for the two most extreme simulated prevalence rates. We left out the highest prevalence results due to division-with-zero difficulties in the multiple threshold scenario.

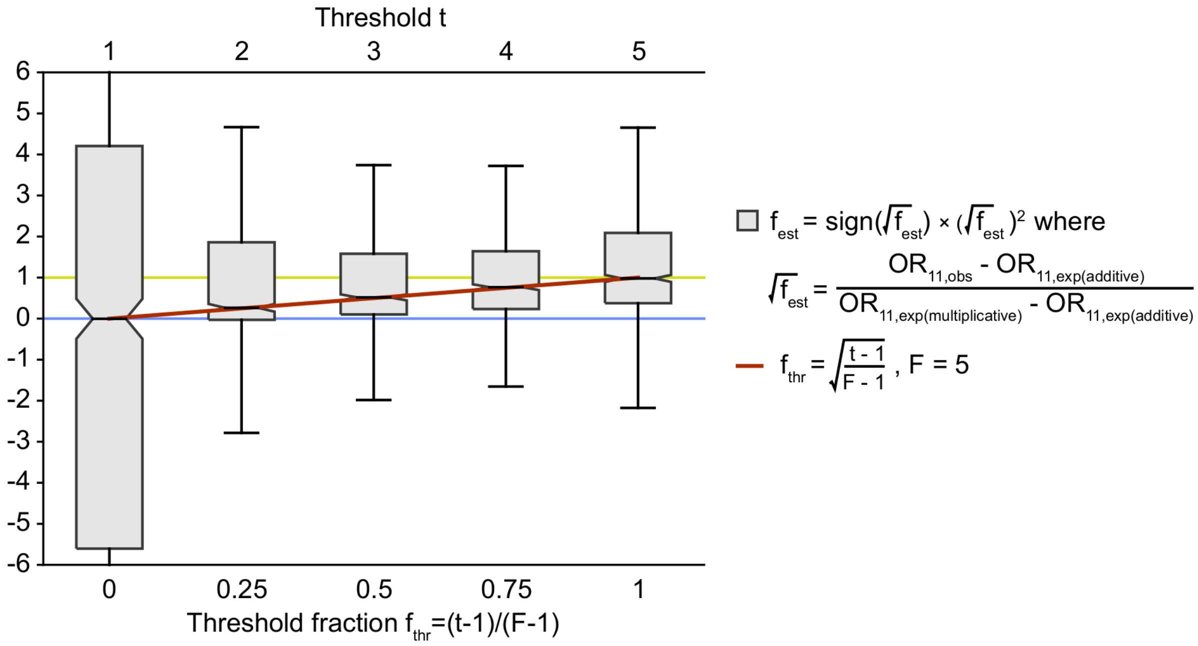


**S2 Fig. Multifactorial threshold model with $f_{est}$ metric.** The y axis is the signed square of the y axis in Fig 2A, otherwise this is the same plot, i.e. based on the scenario in Fig 1C. The red curve is the threshold fraction; this and $f_{est}$ make up the y axis. Notched in the box plots show bootstrapped 95% confidence intervals for the medians. The simulation had 0.5% cases and the rest controls, with a range ±0.04%.

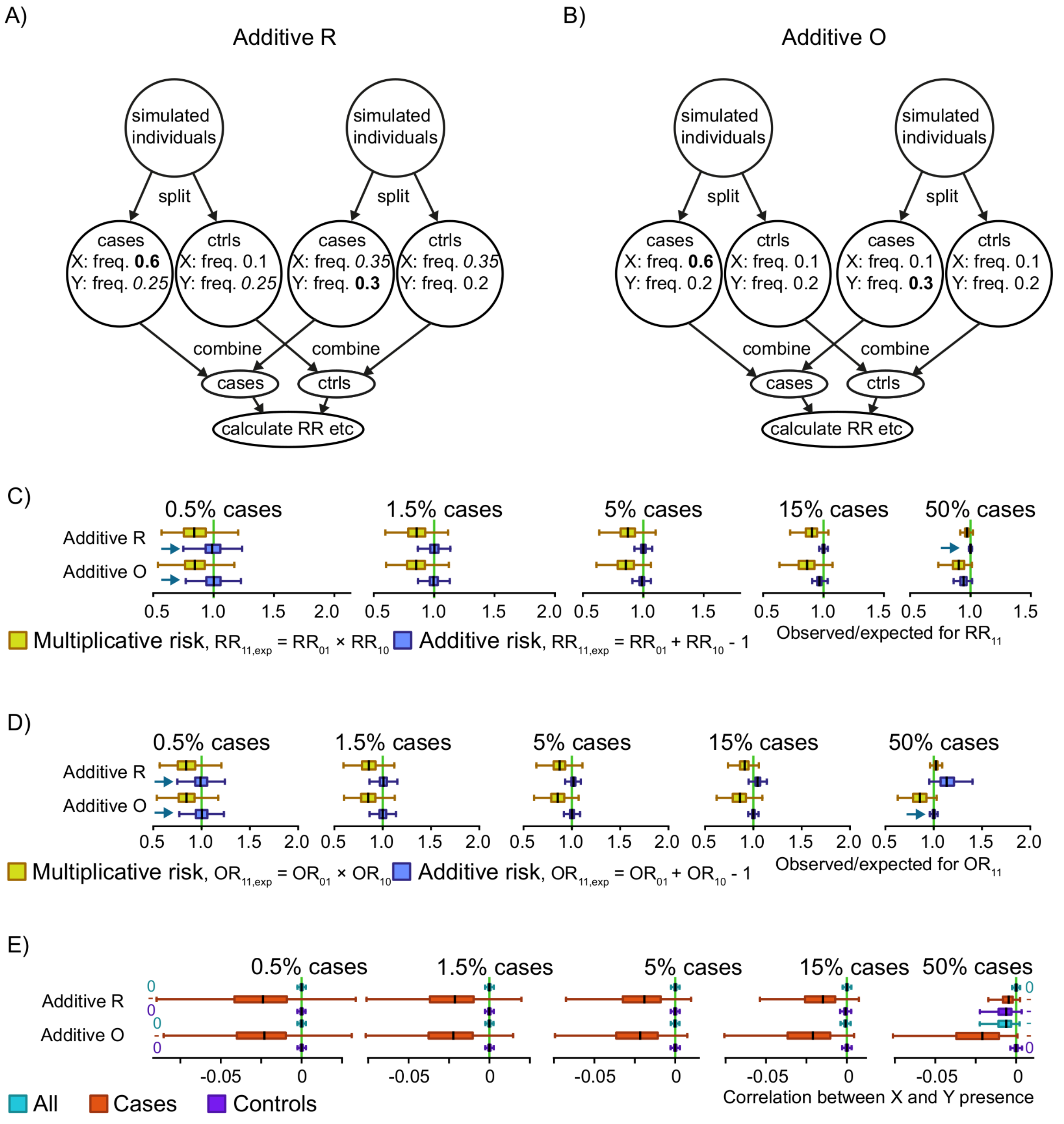


**S3 Fig. Schemes providing additivity.** A-B) Simulation schemes for two schemes intended to provide additive effect. The numbers are example frequencies, and frequencies in bold highlight the higher frequencies of the simulated risk factors (X and Y) associated with disease. For example, “Y: freq. 0.3” means that each simulated individual in the group had a 30% chance of being assigned the risk factor Y. The numbers in italics are the average frequency in the other group of simulated individuals, note that this will depend on how many cases there are to controls, specifically higher_freq · case_fraction + lower_freq · (1 – case_fraction). In each indicated circle the two risk factors have been added in an uncorrelated/independent manner. C-D) The relative risks (C) and odds ratios (D) for double risk ($RR_{11}$ or $OR_{11}$) calculated from the simulation schemes, with boxes summarizing 1,000 simulation runs with different risk factor frequencies. The observed $RR_{11}$ or $OR_{11}$ were compared to the additive and multiplicative combinations of the relative risks for single risk. Boxplots show median and quartiles for the simulations, but extreme values are omitted for clarity. Yellow arrows highlight where the median is visibly close to multiplicativity, while blue arrows do the same for additivity, for the two most extreme simulated prevalence rates. E) Correlation coefficients between the risk factors X and Y. The relevant signal in each case is whether the median is negative, zero or positive, highlighted with a -, 0 or + symbol for the two most extreme simulated prevalence rates.

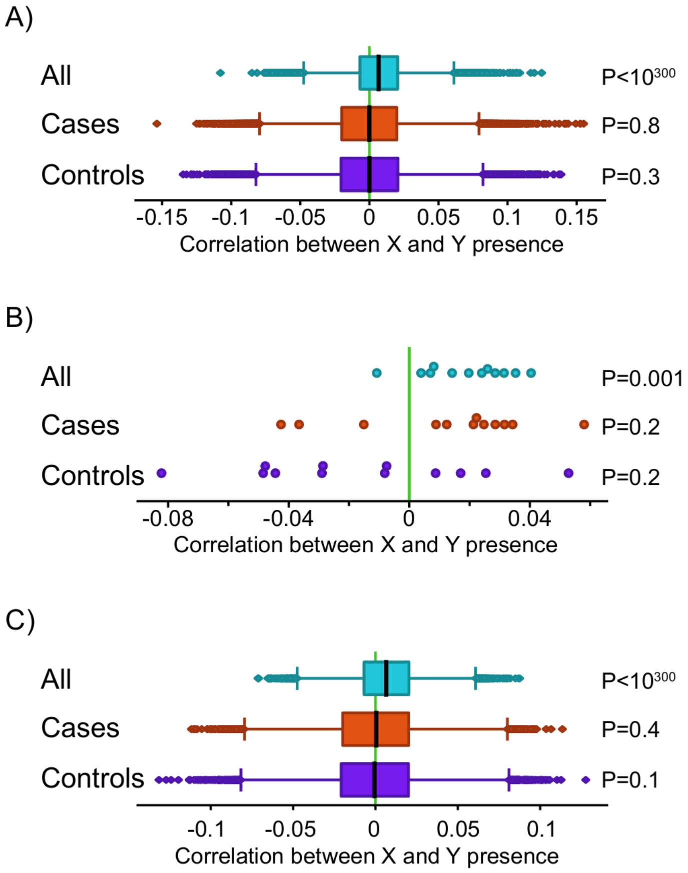


**S4 Fig. Shared epitope correlation relationship.** Correlation between *HLA-DRB1* shared epitope (calculated as codominant) and other SNPs (calculated as dominant) in EIRA data within cases, control or all samples, for all non-HLA SNPs (A, C) or only known non-HLA risk SNPs (B). *P*-values come from 1-sample t-tests against zero. Regressing out sex and the first ten principal components gave essentially the same result (C) as without this step (A, B).

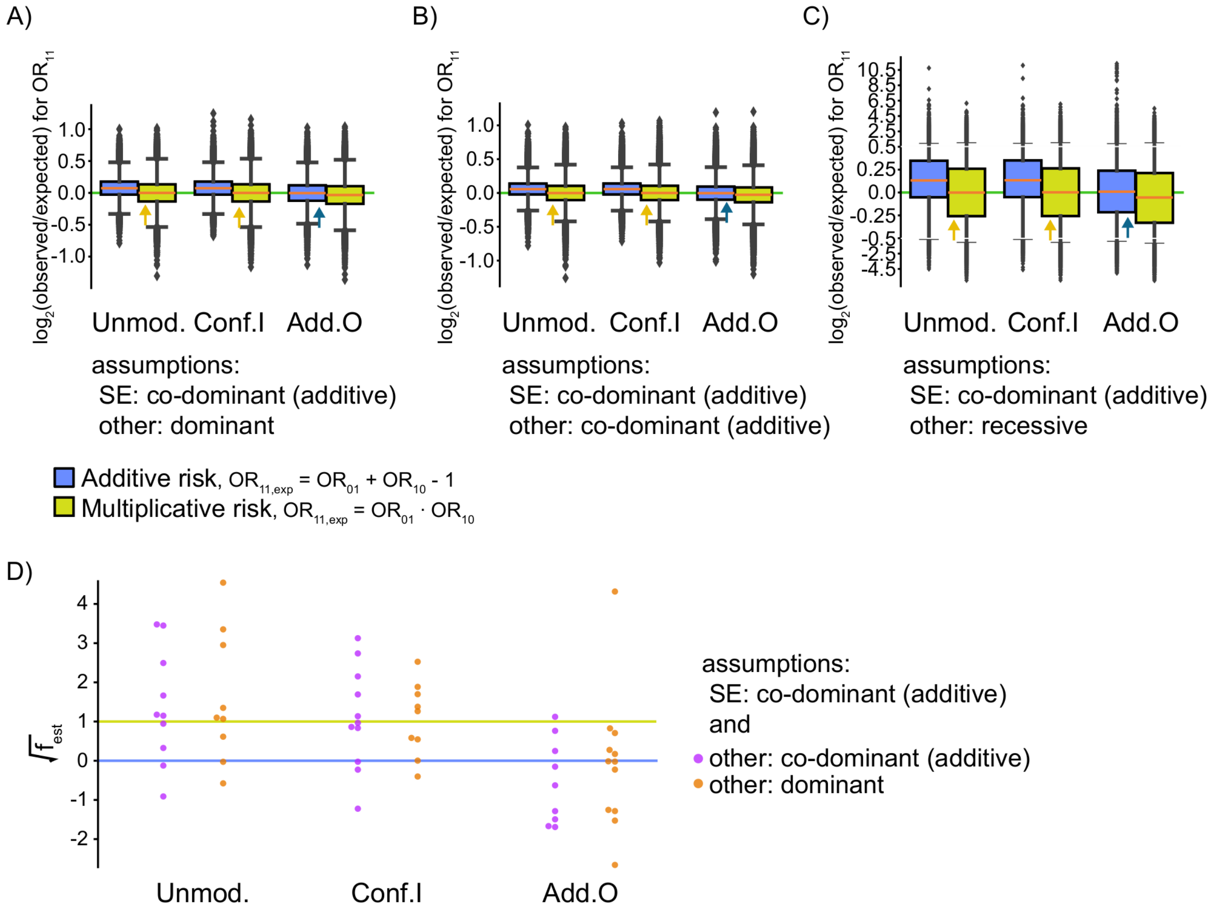


**S5 Fig. Comparison of EIRA double risk odds ratios to two types of randomizations.** A-C) SE is *HLA-DRB1* shared epitope, “other” are non-HLA risk SNPs for ACPA-positive RA. Unmod. is the EIRA data as it is, Conf.I shuffles, for each locus independently, within cases and shuffling within groups to match the independence part of the confounder scenario of Fig 3A, Add.O assigns individuals randomly into two groups and then in one group samples SE from controls and in the other group samples the other SNP from controls, to match S2B Fig. Conf.I preserves one-locus odds ratios and allele frequencies, whereas Add.O unfortunately dampens odds ratios (data not shown), meaning that while Conf.I is a good multiplicative null model, Add.O could use a better replacement. Unmod. and Conf.I have similar variability ($P$=0.04-0.8, Levene’s test, excluding (C) where calculations failed (NaN)). The middle of the scale for (C) is magnified to ease viewing. Blue arrows highlight where the median is visibly close to multiplicativity, while brown arrows do the same for additivity. D) Genes from Figure 4C-D, in varying numbers due to division-by-zero errors, with effect scaled between additive (blue line) and multiplicative (yellow line), as in Fig 2A.
